# Improving the usability and comprehensiveness of microbial databases

**DOI:** 10.1101/497867

**Authors:** Caitlin Loeffler, Aaron Karlsberg, Lana S. Martin, Eleazar Eskin, David Koslicki, Serghei Mangul

**Author notes:** These authors contributed equally to this work (Caitlin Loeffler, Aaron Karlsberg).

## Abstract

Metagenomics studies leverage genomic reference databases to generate discoveries in basic science and translational research. However, current microbial studies use disparate reference databases that lack consistent standards of specimen inclusion, data preparation, taxon labelling and accessibility, hindering their quality and comprehensiveness, and calling for the establishment of recommendations for reference genome database assembly. Here, we analyze existing fungal and bacterial databases and discuss guidelines for the development of a master reference database that promises to improve the quality and quantity of omics research.

## Main Text

High-throughput sequencing has revolutionized microbiome research by enabling the detection of thousands of microbial genomes directly from their host environments [1]. This approach, known as metagenomics, is capable of capturing the complex interactions that take place between thousands of different microbial organisms in their natural habitats. Metagenomic methods rely on comparisons of a sampled genome to multiple reference genomes. Metagenomics is more expensive to perform than traditional, culture-based taxonomic identification techniques, but today’s metagenomic methods can produce a more comprehensive reconstruction of microbial genomes [2]. Emerging technologies for identifying and analysing microbial genomes can provide valuable insights into the interactions between human microbiomes and medicines. However, the current ad hoc practice of storing reference genomes in multiple, disparate reference databases challenges the accuracy and comprehensiveness of future microbial metagenomics studies.

Metagenomic studies isolate DNA found in a sample of various environments, compare the sampled genomes (represented as a set of reads) to verified reference genomes, and identify the organism from which the reads originated. Ideally, a metagenomic study uses a reference database that contains all known genomic references. Today’s researcher can choose from many different genomic reference databases that contain verified reference genomes, but these databases lack a universal standard of specimen inclusion, data preparation, taxon labelling, and accessibility.

Several limitations in genomic sequencing present unique challenges to accurately assembling reference genomes and compile them into comprehensive databases. Notably, reference genomes can exist in various stages of completion. Typically, reads are assembled into larger sequences which represent complete or fragmented microbial genomes. Fragmented assemblies are usually represented as a set of contigs, which are typically contiguous DNA fragments corresponding to unlocalized segments of microbial genomes. Given sufficient data, contigs can be further assembled into scaffolds that represent larger portions of individual chromosomes but gaps (consisting of a possibly unknown number of unknown nucleotides) can remain. Most reference genomes are in different stages of completeness, with portions of even the human genome remaining unknown (in particular, the centromere and telomere regions).

In addition, the location of possibly incomplete reference genomes on taxonomic or phylogenetic trees can be contentious. Metagenomics researchers must take into account discrepancies in the types of taxa included in each reference genome database, as well as differences in how the genomes are constructed, identified, and made available for distribution.

The future of metagenomics research would benefit from a standardized, comprehensive approach to reference genome database development. To begin assembling a set of recommendations for reference genome database construction, we assessed the concordance and usability of available reference databases for microbial genomics. Our study considered the concordance of microbial species and genera across four fungal reference databases (Ensembl [3], RefSeq [4], JGI’s 1000 fungal genomes project (JGI 1K) [5], and FungiDB [6]) and three bacterial reference databases (Ensembl [3], RefSeq [4], and PATRIC [7]). We compared the microbial taxa in each of the databases using NCBI’s universal taxonomic identifiers (hereafter referred to as taxIDs) at the ranks of species and genus (Additional File 1). Strains were not included in this analysis as studied databases contained multiple instances where a reference was counted as a strain in one database yet was labelled an isolate in NCBI; in such cases, the reference was not yet assigned a strain-level NCBI taxID. This discrepancy made comparison of strain comprehensiveness among databases impossible to calculate and demonstrates the importance of developing a standardized taxonomic naming system to be shared between databases [8].

Our comparison of four major fungal and three major bacterial genome databases reveals substantial discrepancies across databases in the presence of microbial references at taxonomic levels below the family rank. In other words, a researcher’s selection of one particular reference database could substantially impact the number and types of unique microbial taxa identified in a study.

Calculating the coverage of each fungal reference genome database shows that a researcher using the largest—and most comprehensive— reference database would only find identification for 80% of the possible 1405 fungal species (Fig. 1a) and 95% of the possible 42,337 bacterial species (Fig. 1b). For genera, a researcher using the largest—and most comprehensive— fungal reference database would only find identification for 89% of the total 786 genera covered by all four fungal databases (Fig. 1c) and 94% of the total 3371 genera covered by all three bacterial databases (Fig. 1d)

**Figure 1.**
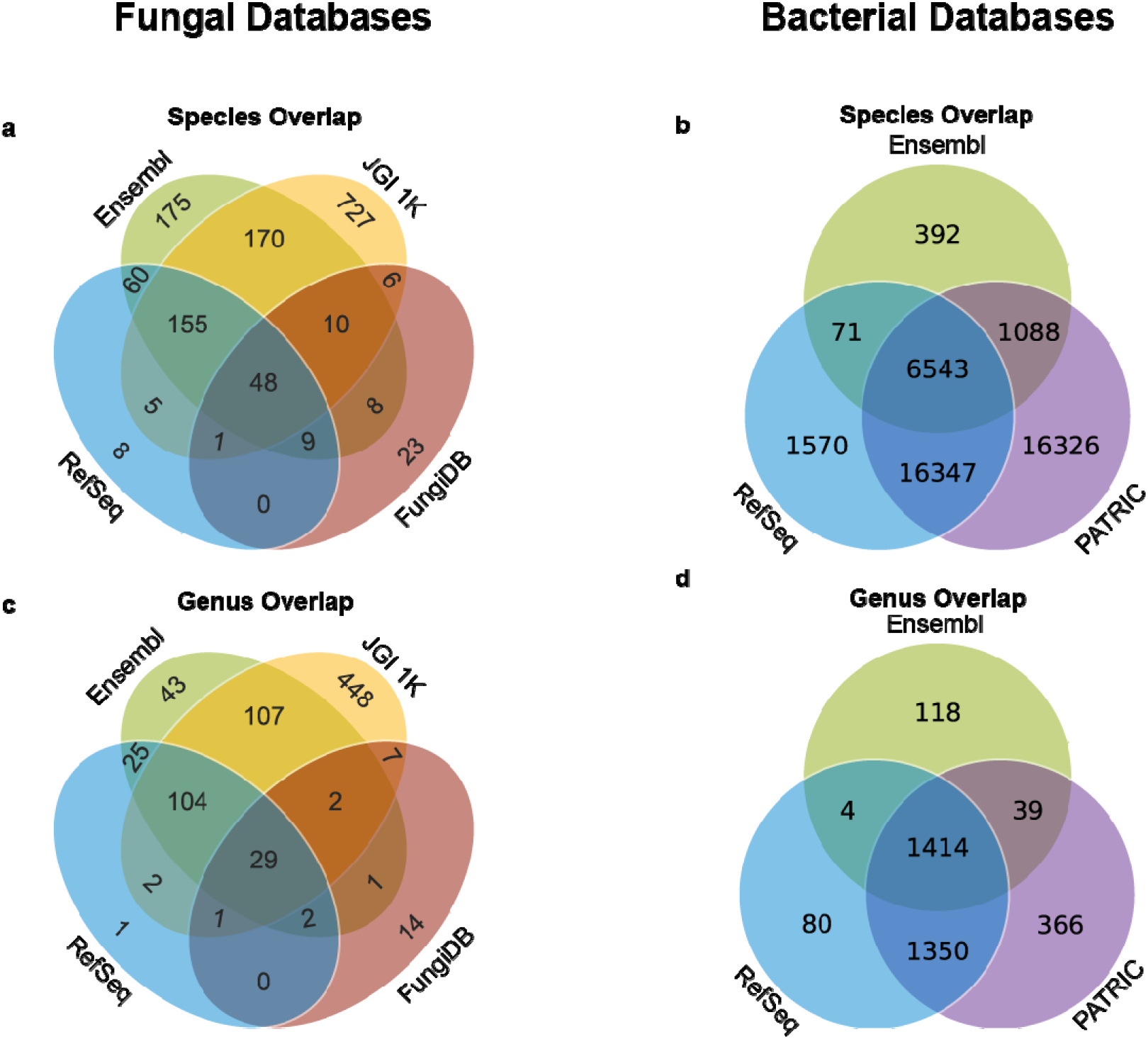
Consensus of fungal and bacterial genome representation across multiple reference databases. (a) In total, there are 1405 unique species represented across the four fungal databases. Of these, 48 species are represented in all four databases. There are a total of 175 species found where strictly three databases overlap and 189 species where strictly two databases overlap. A total of 993 unique fungal species cannot be found in any overlaps. (b) In total, there are 42337 unique species represented across the three bacterial databases. Of these, 6543 species are represented in all three databases, and 17506 total species are found where strictly two databases overlap. A total of 18288 unique bacterial species cannot be found in any overlaps. (c) In total, there are 786 unique genera represented across the four fungal databases. Of these, 29 genera are represented in all four databases. There are a total of 109 genera found where strictly three databases overlap and 142 genera where strictly two databases overlap. A total of 506 unique fungal genera cannot be found in any overlaps. (d) In total, there are 2214 unique genera represented across the three bacterial databases. Of these, 76 genera are represented in all three databases, and 1149 total genera are found where strictly two databases overlap. A total of 989 unique bacterial genera cannot be found in any overlaps.

Only a relatively small percentage of species are represented as complete genomes; calculating the percentage of fungal species per reference database reveals that 16% of species are represented as complete fungal genomes in Ensembl, 2% in JGI 1K, 14% in RefSeq, and 13% in FungiDB. Conversely, our study shows that the percentage of species represented as contigs are relatively high: 81% in RefSeq, 98% in JGI 1K, 80% in Ensembl, and 81% in FungiDB. Remaining genomes are comprised of contigs or a mixture of chromosomes and contigs (Fig. 2a). In addition, we found that complete reference genomes for fungi taxa were not consistently present in studied fungal reference databases. In total, there are 53 unique species represented across the four fungal databases that are complete genomes. Of these, only 13% are represented in all four databases (Additional File: Fig. S1).

**Figure 2.**
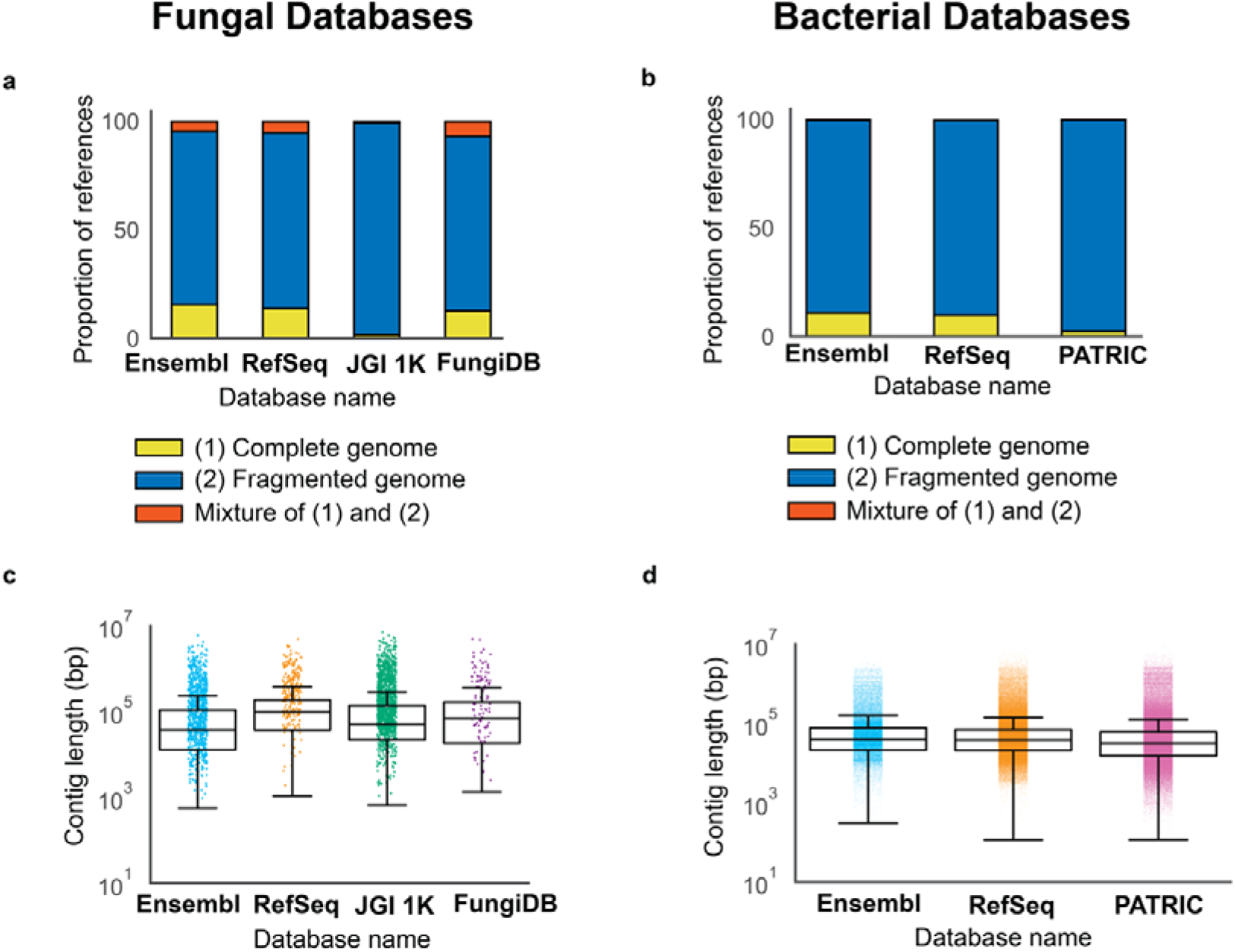
Fungal and bacterial genome composition across multiple reference databases. (a) Percentage of references per fungal database available as complete genomes (yellow), fragmented genomes (i.e., set of contigs) (blue), and a mixture of full chromosomes and contigs (red). (b) Percentage of species per bacterial database available as complete genomes (yellow), fragmented genomes (i.e., set of contigs) (blue), and a mixture of full chromosomes and contigs (red). (c) Length distribution of the fungal genomes (contigs) across the databases. The contig mean lengths for each fungal database are 322 thousand bp (Ensembl), 513 thousand bp (RefSeq), 426 thousand bp (JGI 1K), and 548K (FungiDB). (d) Length distribution of the bacterial genomes (contigs) across the databases. The contig mean lengths for each bacterial database are 149 thousand bp (Ensembl), 119 thousand bp (RefSeq), and 107 thousand bp (PATRIC).

We found similar results for bacterial species in the bacterial genome reference databases. Only 11% of bacterial references are represented as complete bacterial genomes in Ensembl, 10% in RefSeq, and 3% in PATRIC. The majority of references are represented as contigs in Ensembl (89%), RefSeq (90%), and PATRIC (97%). All three bacterial genome reference databases have <1% of references containing a mix of contigs and chromosomes (Fig. 2b).

Of the 80 - 90% of the references in each database represented as fragmented genomes, we considered the length distributions of the sequences provided. The length distributions for contigs are relatively similar across all four fungal databases (Fig. 2c). The length distributions for contigs are relatively similar across the three bacterial databases we studied (Fig. 2d). The mean contig length is shorter in bacterial reference databases than in fungal reference databases.

The completeness of a reference database is always subject to limitations imposed by the project’s funding or scope. As one example of the latter, the JGI 1K database contains many novel and previously unpublished genomes. The introductory text of the JGI database indicates that, for this reason, it is not designed to be used in metagenomics studies [5]. However, such a large database of novel references may be a top choice for metagenomics researchers who want to learn as much as they can about their samples. Of the four fungal reference databases analyzed in this study, JGI 1K is the largest, covering 89% of fungal genera and 80% of fungal species. Ensembl, the second largest of the four databases, only covers 45% of fungal species and 41% of fungal genera.

In some cases, a more complete database may hinder analytical methods. Due to limitations in metagenomic analysis pipelines, reference databases containing species whose genomes are remarkably similar often prevent identification at the species level [9].

Even taking these limitations into account, researchers would benefit from a universal approach to constructing comprehensive microbial genomic reference databases. Since the ideal reference database containing all the reference genomes for all known samples does not yet exist, researchers are potentially failing to identify key organisms within their samples. The first consideration of a master reference database would be developing a standardized approach to assembling and presenting data from existing reference databases. A systematic approach to constructing reference databases, when adopted by the scientific community, would help improve microbial coverage in newly developed metagenomic analysis tools.

One approach to developing a comprehensive database of complete genomic references is to combine all existing reference databases into one master set—a complex, time-consuming task. With this approach, references unique to one database could simply be added to a master set. However, a reference that is found in more than one database presents several problems. Multiple references may be assigned the same taxID, yet these references may contain differing genomic information. For example, references comprised of contigs could cover different segments of a given gene. Selecting both unedited contig-based reference genomes would unnecessarily extend the run time of a comparison algorithm utilizing the master set. On the other hand, eliminating one reference would ignore entire segments of the genome represented in the discarded contigs. In such cases, the database developer needs a consistent method for selecting one of the references to include in the master set.

An alternative approach would be to develop an open source computational method that continuously merges any number of disjointed microbial reference databases as new sequences become available. The sequencing and storing of microbial species in multiple repositories present an opportunity to improve sequence quality through an approach based on alignment and consensus. An open source format would encourage computational developers to contribute to the reference database by engineering support for the integration of other, lesser known, reference sequence repositories.

Another potential strategy is to eliminate discrepancies between databases. This will require the development of a communication protocol that allows databases to share information and complement each other in real time. Such a communication protocol could eventually enable an assembly of a comprehensive ‘virtual’ database, which essentially represent a consensus across databases. Several technical issues may pose difficulties in implementing such an approach. For example, the proposed approach needs to be capable of resolving the conflicts between the databases, such as when references are represented by different contigs across databases.

We would also like to mention that, just like reference databases for genetic data, the reference databases for taxonomies also have restricted overlap [8]. For this present study, we were able to use NCBI Universal Taxonomic IDs (taxIDs) to measure species and genus reference congruence across the databases since NCBI taxIDs were used by each database we studied. Hence, database discrepancies only existed due to presence or absence of organisms in the reference database, not due to taxonomic ambiguities. However, there exist many such universal taxonomic systems which may overlap very little and where there may not exist a mapping system to convert from one taxonomy to another. Further, even though we were able to identify species and genus across databases by NCBI taxIDs, this did not extend to strains as NCBI does not universally assign taxonomic identifiers to strains. The master database for reference genomes, will, therefore, also need to utilize a master database for taxonomy. For example, one possible master taxonomic database may be the OpenTree taxonomy [8].

A second consideration of a master reference database is usability. Bioinformatics is an interdisciplinary field comprised of researchers with varied backgrounds—from computer science to biology. In order to maximize potential use by both skilled and novice computational users, this complete database would need an intuitive user interface.

The four fungal and three bacterial databases analysed in this study presented challenges to data access and manipulation. For example, the fungal JGI 1K asks the user to select the genomes of interest from a picture of the fungal tree of life, which can be unintuitive to many researchers. Adequate user support would also increase the usability of a comprehensive reference database; at the time of our study, Ensembl did not publish any contact information on their webpage.

Several reference databases highlight features that should be implemented in a master reference database. The interface for FungiDB, which is more intuitive, simply asks the user to select data as though shopping online. To download all organisms, one only had to hover over “About FungiDB”, click “Organisms” under “------- Data in FungiDB”, click “add to basket”. Once all the organisms are placed in the basket, it is possible to customize an annotation table containing download links for all references within the basket. While downloading data from NCBI RefSeq can be challenging, once the user knows to select “Assembly” in the dropdown menu on the home page and type “Fungi” into the search bar, the filtering process becomes more intuitive. The “shopping basket” method is not efficient for downloading bacterial references, however, as there are over 200 thousand references to handle. A better approach would be to allow the user to download references from the NCBI FTP site, which requires knowledge of the command line and may not be usable by researchers lacking a computational background.

A third consideration of a master reference database is maintenance support and archival stability. Maintaining a master reference sequence database would carry a substantial cost in terms of computational power and storage. An open source, continuous assembly approach would depend on support from an institution, governing body, or a global consortium.

The Pathosystems Resource Integration Center (PATRIC) online bacterial reference database can be used as a gold standard for database website design. In PATRIC, all genomes for the selected taxa are present, and the filtering is intuitive. One drawback to the PATRIC website is the current protocol for downloading genomes; the best way to transfer data between servers is to generate a list of genome_ids in the command line for the genomes of interest, then recursively call “wget” on each genome. Any researcher not familiar with the command line needs to download the data directly from the PATRIC website; this method is not allowed for bulk downloads. A more efficient alternative method for bulk downloading reference data without using the command line would be to provide an option to utilize a data transfer service (such as Globus), which PATRIC does not currently use.

Our study indicates that the current approach to developing genomic reference databases for fungal and bacterial species are not meeting the needs of metagenomics research. As the resolution of metagenomic data increases, researchers will have more need for tools that precisely identify the taxonomy of DNA derived from samples. We believe that a systematic approach to developing a centralized master reference database will increase coverage and dramatically improve the quality and quantity of - omics research.

**Declarations**

## Acknowledgements

Not Applicable

## Availability of Data and Materials

The data supporting the conclusions of this article, including the species and genera names, are available at: https://github.com/Mangul-Lab-USC/db.microbiome.

## Author contributions

C.L. and A.K. performed the analysis. C.L., L.M., D.K., and S.M. wrote the manuscript. S.M. conceived of the presented idea. E.E. supervised the project. All authors read and approved the final manuscript.

## Authors’ information

C.L.; Twitter @LoefflerCaitlin

A.K.; Twitter @A4RONK

L.S.M.

E.E.

D.K.; Twitter @DavidKoslicki

S.M.; Twitter @serghei_mangul

## Tables

Not Applicable

## Additional Files

Additional File 1; Supplementary Material; Detailed methods; Additional File 1.docx

Additional File 1 Figure S1; Overlap of complete genomes; A venn diagram showing the distribution of complete chromosome coverage for the four fungal databases; Additional File 1 Figure S1.png

## References

1. Sharpton TJ. An introduction to the analysis of shotgun metagenomic data. Front Plant Sci. 2014 Jun 16;5:209.

2. Hilton SK, Castro-Nallar E, Pérez-Losada M, Toma I, McCaffrey TA, Hoffman EP, Siegel MO, Simon GL, Johnson WE, Crandall KA. Metataxonomic and metagenomic approaches vs. culture-based techniques for clinical pathology. Front Microbiol. 2016 Apr 7;7:484.

3. Kersey PJ, Allen JE, Allot A, Barba M, Boddu S, Bolt BJ, Carvalho-Silva D, Christensen M, Davis P, Grabmueller C, Kumar N. Ensembl Genomes 2018: an integrated omics infrastructure for non-vertebrate species. Nucleic Acids Res. 2018 Jan 4;46(D1):D802–8.

4. O’Leary NA, Wright MW, Brister JR, Ciufo S, Haddad D, McVeigh R, Rajput B, Robbertse B, Smith-White B, Ako-Adjei D, Astashyn A. Reference sequence (RefSeq) database at NCBI: current status, taxonomic expansion, and functional annotation. Nucleic Acids Res. 2016 Jan 4;44(D1):D733–45.

5. Nordberg H, Cantor M, Dusheyko S, Hua S, Poliakov A, Shabalov I, Smirnova T, Grigoriev IV, Dubchak I. The genome portal of the Department of Energy Joint Genome Institute: 2014 updates. Nucleic Acids Res. 2014 Jan 1;42(D1):D26–31.

6. Basenko EY, Pulman JA, Shanmugasundram A, Harb OS, Crouch K, Starns D, Warrenfeltz S, Aurrecoechea C, Stoeckert CJ, Kissinger JC, Roos DS. FungiDB: an integrated bioinformatic resource for fungi and oomycetes. J Fungi. 2018 Mar;4(1):39.

7. Wattam AR, Davis JJ, Assaf R, Boisvert S, Brettin T, Bun C, Conrad N, Dietrich EM, Disz T, Gabbard JL, Gerdes S. Improvements to PATRIC, the all-bacterial bioinformatics database and analysis resource center. Nucleic Acids Res. 2017 Jan 4;45(D1):D535–42.

8. Hinchliff CE, Smith SA, Allman JF, Burleigh JG, Chaudhary R, Coghill LM, Crandall KA, Deng J, Drew BT, Gazis R, Gude K. Synthesis of phylogeny and taxonomy into a comprehensive tree of life. Proc Natl Acad Sci U S A. 2015 Oct 13;112(41):12764–9.

9. Nasko DJ, Koren S, Phillippy AM, Treangen TJ. RefSeq database growth influences the accuracy of k-mer-based lowest common ancestor species identification. Genome Biol. 2018 Dec;19(1):1–0.

