## Supplementary Materials for "Improving the usability and comprehensiveness of microbial databases"

{Additional File 1}

Part I: Downloading the databases

Our study considered fungal species and genera across four reference databases:

- JGI 1000 Fungal Genomes Database (JGI 1K), https://genome.jgi.doe.gov/programs/fungi/index.jsf
- Ensembl, http://fungi.ensembl.org/index.html
- RefSeq, https://www.ncbi.nlm.nih.gov/
- FungiDB, http://fungidb.org/fungidb/

Our study also considered bacterial species and genera across three reference databases:

- Ensembl, http://bacteria.ensembl.org/index.html
- RefSeq, https://www.ncbi.nlm.nih.gov/
- PATRIC, https://www.patricbrc.org/

Each reference database had a different process for downloading reference genomes. For fungi:

- **JGI 1K Fungal Genomes Database.** We locally downloaded the assembled masked fungal reference database, which appeared in the “download” section of the website as a zipped file. The unzipped file yielded 1265 directories. Each directory represented one species or, when strain information was available, one strain. The contents of each directory included a zipped FASTA file (in two directories there were two such files) that contained the genetic reference information. Plasmid and mitochondrial sequences were available in separate files.
- **Ensembl.** We called wget recursively on all FTP files in release 44 ending in ‘.dna.toplevel.fa.gz’.
- **RefSeq.** We downloaded the FASTA files locally after filtering the NCBI assembly “Fungi” results for latest RefSeq references.
- **FungiDB.** Once all the fungal reference genomes were placed into the online basket, we downloaded the links corresponding to FASTA files. We then created a bash file that called “wget” on each link.

For bacteria:

- **Ensembl.** We ran a simple one line script accessing the FTP site
- **RefSeq.** We ran a recursive wget function that downloaded all the bacterial fasta files.
- **PATRIC.** The FTP site does not group genomes by lineage, and bulk downloads from the website itself is limited to sets of 10 thousand references at a time. A list of genome ids was generated in a text file which was then accessed by a wget loop to download each individual genome of interest.

Part II: Standardizing the taxonomy across the fungal and bacterial reference databases

In order to standardize the taxonomy across all four fungal and three bacterial reference databases, provided NCBI universal taxIDs were used in place of scientific names. TaxIDs are given at each taxonomic level and, therefore, can be ranked from Superkingdom to Strain levels. Only species- and genus-level taxIDs were used to quantify the consensus of fungal and bacterial genome representation across the databases. We did not analyze the consensus of strain-level taxIDs. We used the Ete3 module to assign a species-level taxID when the database provided a strain-level taxonomic identification, and to obtain a genus-level taxID for all reference genomes. We encountered multiple genomes that had not been assigned a genus and lacked a genus-level taxID; in the data, such reference genomes indicated ‘no rank’ where the genus-level taxID would have appeared. In such cases, we used the unranked taxID as the genus taxID.

As with the procedures for downloading reference genomes, the processes for obtaining corresponding taxIDs for each file were different for each of the databases.

- **JGI 1K Fungal Genomes Database.** We followed the six-step process for obtaining a Microsoft Excel document with the taxIDs. First, we created an advanced search that produced a “reports” button where the user can download the taxID information via an Excel spreadsheet. Second, we prepared a Python script to automatically match filenames with corresponding taxIDs.
- **Ensembl.** The FTP server (ftp://ftp.ensemblgenomes.org/pub/fungi/release-44/) offers a file named “species_EnsemblFungi.txt,” a mapping file that shows which Ensembl files match to corresponding taxIDs. We ran a Python script that used the mapping file and a list of Ensembl file names to assign taxIDs.
- **RefSeq.** Similar to Ensembl, we used a mapping file to match taxIDs to accession numbers listed in the first header of each reference FASTA file.
- **FungiDB.** After downloading all the reference species, a csv file could be custom generated by clicking the “download” link, selecting “choose columns”, and checking the “NCBI taxon ID” box under “taxonomy”.

For the bacterial databases we isolated a list of taxIDs present in a given bacterial database and did not match taxIDs to filename.

Part III: Classify reference genomes as complete or fragmented

In order to determine the assembly level (e.g., scaffolds, contigs, fully assembled chromosomes) and identity of extra genetic material (e.g., mitochondrial and plasmid sequences) for each reference genome, we searched the headers of each reference FASTA file for predetermined patterns and words. We used the substrings “chr”, “complete”, and “NC_” to identify sequences that had been marked as complete genomes. The key substrings “contig”, “scaffold”, “partial”, “supercont”, “unitig_”, “NW_”, and “NT_” were used to identify sequences that had been marked as fragmented genomes.

When identifying the extra genetic material, we used the substrings “mitochondri” and “mt dna” to identify sequences that had been marked as mitochondrial references. Finally, we used the word “plasmid” to identify sequences that had been marked as plasmids.

Part IV: Compare the species and genera across the reference databases

In order to generate statistical data for cross-database reference comparison, we extracted individual sequence attributes from each reference FASTA file. We stored these attributes in a structured query language relational database management system (SQL RDBM). Attributes extracted from each fungal reference sequence included database name; species-level taxID; genus-level taxID; species name; genus name; a flag indicating reference composition (e.g., chromosomes, contigs or mixture of both); a flag indicating if the reference contains mitochondrial and plasmid DNAs.

For each reference, we also recorded the length of contigs and chromosomes. Individual files could have more than one sequence classification depending on the contents of the DNA sequences. The data for sequence composition contained the number of sequences for a given sequence classification that existed within each file. We also stored within each file the average, minimum, and maximum sequence lengths for each sequence classification.

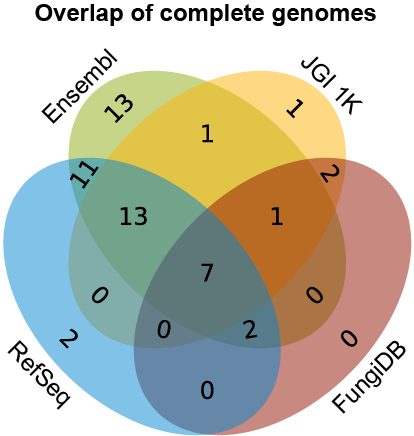

**Figure S1. Overlap of species level references containing only complete chromosomes.** In total, 53 unique species references contain only complete chromosomes represented across the four fungal databases. Of these, seven species are represented in all four databases. There is a total of 16 species found where strictly three databases overlap and 13 species where strictly two databases overlap. A total of 17 unique fungal species cannot be found in any overlaps.
