## Supplementary figures and images for "Improving the usability and comprehensiveness of microbial databases"

### Figure S1

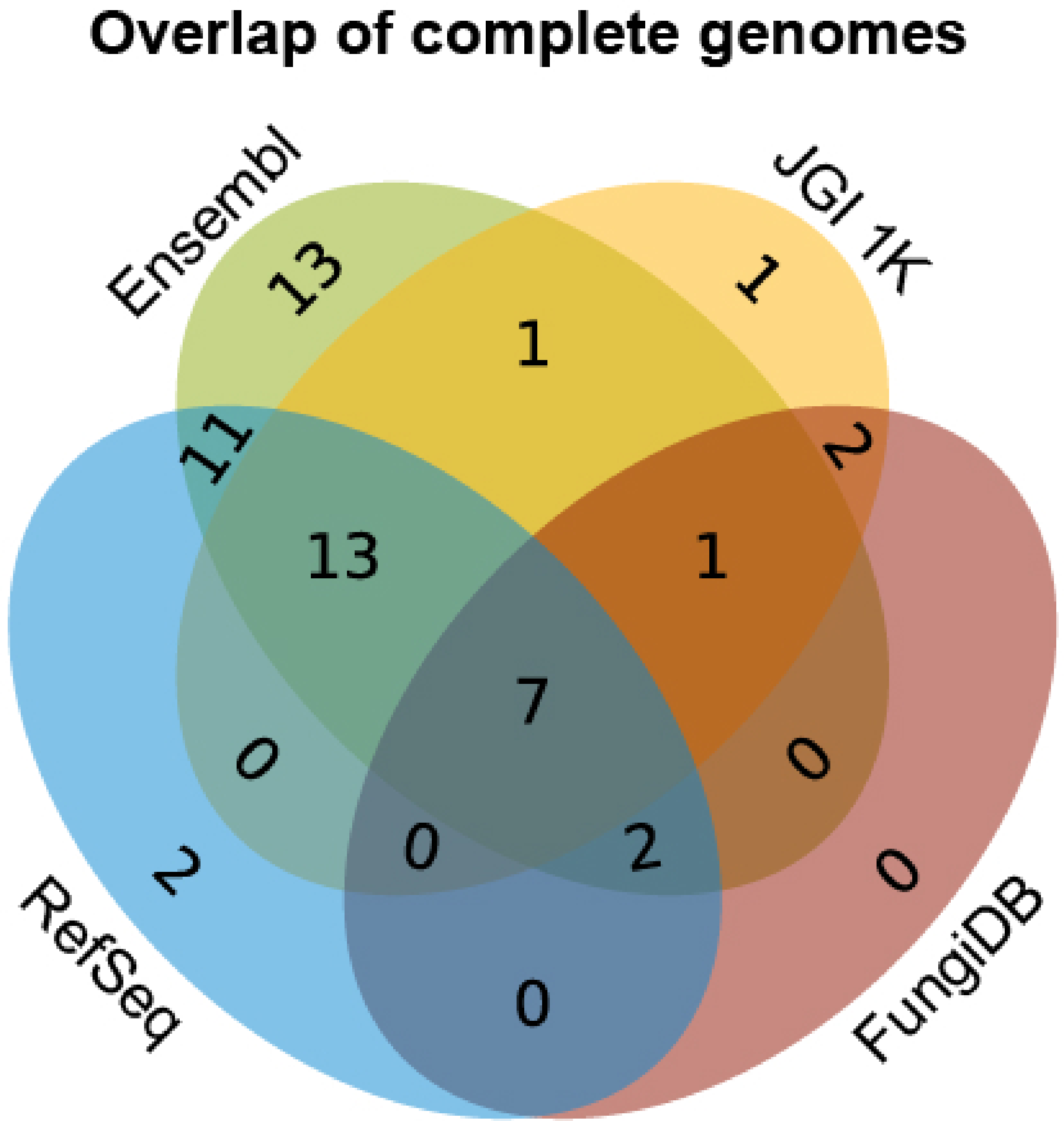
